# A Modular Workflow for Model Building, Analysis, and Parameter Estimation in Systems Biology and Neuroscience

**DOI:** 10.1101/2020.11.17.385203

**Authors:** João P.G. Santos, Kadri Pajo, Daniel Trpevski, Andrey Stepaniuk, Olivia Eriksson, Anu G. Nair, Daniel Keller, Jeanette Hellgren Kotaleski, Andrei Kramer

## Abstract

Neuroscience incorporates knowledge from a range of scales, from molecular dynamics to neural networks. Modeling is a valuable tool in understanding processes at a single scale or the interactions between two adjacent scales and researchers use a variety of different software tools in the model building and analysis process. While systems biology is among the more standardized fields, conversion between different model formats and interoperability between various tools is still somewhat problematic. To offer our take on tackling these shortcomings and by keeping in mind the FAIR (findability, accessibility, interoperability, reusability) data principles, we have developed a workflow for building and analyzing biochemical pathway models, using pre-existing tools that could be utilized for the storage and refinement of models in all phases of development. We have chosen the SBtab format which allows the storage of biochemical models and associated data in a single file and provides a human readable set of syntax rules. Next, we implemented custom-made MATLAB^®^ scripts to perform parameter estimation and global sensitivity analysis used in model refinement. Additionally, we have developed a web-based application for biochemical models that allows simulations with either a network free solver or stochastic solvers and incorporating geometry. Finally, we illustrate convertibility and use of a biochemical model in a biophysically detailed single neuron model by running multiscale simulations in NEURON. Using this workflow, we can simulate the same model in three different simulators, with a smooth conversion between the different model formats, enhancing the characterization of different aspects of the model.

**Information Sharing Statement:** Both the source code and documentation of the Subcellular Workflow are available at https://github.com/jpgsantos/Subcellular_Workflow and licensed under GNU General Public License v3.0. The model is stored in the SBtab format (Lubitz et al. 2016). Model reduction, parameter estimation and global sensitivity analysis tools are written in MATLAB^®^ (RRID:SCR_001622) and require the SimBiology^®^ toolbox. Conversion script to VFGEN (Weckesser 2008), MOD and SBML (RRID:SCR_007422) is written in R (RRID:SCR_001905). Conversion to SBML requires the use of libSBML (RRID:SCR_014134). Validations are run in COPASI (RRID:SCR_014260; Hoops et al. 2006), NEURON (RRID:SCR_005393; Hines and Carnevale 1997) and with the subcellular simulation setup application (RRID:SCR_018790; available at https://subcellular.humanbrainproject.eu/model/simulations) that uses a spatial solver provided by STEPS (RRID:SCR_008742; Hepburn et al. 2012) and network-free solver NFsim (available at http://michaelsneddon.net/nfsim/). The medium spiny neuron model (Lindroos et al. 2018) used in NEURON simulations is available in ModelDB database (RRID:SCR_007271) with access code 237653. The FindSim use case model is available in https://github.com/BhallaLab/FindSim (Viswan et al. 2018).

## Introduction

Computational systems biology is a data-driven field concerned with building models of biological systems. Methods from systems biology have proven valuable in neuroscience, particularly when studying the composition of synapses, molecular mechanisms of plasticity, learning and various other neuronal processes (Bhalla and Iyengar 1999; Hellgren Kotaleski and Blackwell 2010; Li et al. 2012). A wide variety of different software and toolboxes, each with their own strengths and weaknesses, are available within the field. This diversity, however, can obstruct model reuse as interoperability between the different software packages and the convertibility between various file types is only solved in part. Interoperability can either mean that the model built in one simulator can be run in another or that both simulators interoperate at run-time either at the same or different scales (Cannon et al. 2014). The former is addressed by standardizing model descriptions, and in systems biology by the standard machine-readable model formats as XML-based SBML (Systems Biology Markup Language; Hucka et al. 2003) and CellML (Hedley et al. 2001), and human readable format such as SBtab (Lubitz et al. 2016). An analogous model description language for neurons and networks is the NeuroML (Neural Open Markup Language; Gleeson et al. 2010).

We start by providing examples of the available systems biology tools. We then proceed to describe our approach in developing a modular workflow to address some of these interoperability issues and present simulation results of an example use case in various simulators and frameworks, with further examples provided, in supplementary materials. Our workflow starts with a human-readable representation of the model that is easily accessible to everyone and proceeds through various conversions into different simulation environments: MATLAB^®^, COPASI, NEURON, and STEPS. Specifically, we will describe the conversion tools we created for this purpose.

## Examples of Software and Toolboxes used in Systems Biology

No software package is perfectly suited for every task, some have programmable interfaces with scripting languages, like the MATLAB^®^ SimBiology^®^ toolbox, some focus on providing a fixed array of functions that can be run via graphical user interfaces, like COPASI, although it now offers a Python toolbox for scripting (Welsh et al. 2018). Most toolboxes and software packages offer a mixture of the two approaches: fine-grained programmable interface as well as fixed high-level operations. At the extremes of this spectrum are powerful but inflexible high-level software on one side, and complex, hard to learn but very flexible libraries or toolboxes with an API on the other. Some examples of general modeling toolboxes in MATLAB^®^ are the SBPOP/SBToolbox2 (Schmidt and Jirstrand 2006), and the PottersWheel Toolbox (Maiwald and Timmer 2008). For Bayesian parameter estimation, there are the MCMCSTAT toolbox (Haario et al. 2006) in MATLAB^®^, as well as pyABC (Klinger et al. 2018) and pyPESTO (Schälte et al. 2020) in Python, and the standalone Markov Chain Monte Carlo (MCMC) software GNU MCsim (Bois 2009). For global sensitivity analysis, there is the Uncertainpy Python toolbox (Tennøe 2018). For simulations in neuroscience, examples are NEURON (Hines and Carnevale 1997) and STEPS (Hepburn et al. 2012). Both are used for simulations of neurons and can include reaction-diffusion systems and electro-physiology.

These software packages do not all use the same model definition formats. Most have some compatibility with SBML, others use their own formats (e.g. NEURON uses MOD files). In some cases, an SBML file exported from one of these packages cannot be imported into another package without errors; so manual intervention may be required^1^. Given an SBML file, a common task is to translate the contents into code that can be used in model simulations, for ordinary differential equations this is the right-hand side vector field function. There are several tools that facilitate the conversion between formats, e.g. the SBFC (The Systems Biology Format converter; Rodriguez et al. 2016) as well as the more general VFGEN (A Vector Field File Generator; Weckesser 2008).

All toolboxes and software packages have great strengths and short-comings, and each programming language has different sets of (freely) available libraries which makes the development (or use) of numerical methods more or less feasible than in another language. One such example is the R package CDvine for parameter dependency modeling, which has recently been used for modeling probability densities (in parameter spaces) between two MCMC runs (Eriksson et al. 2019). It infers the probability denisty function from a large enough sample, another method to do this is *kernel density estimation*^2^. The CDvine package implements a much more advanced and robust method of density estimation based on vines and copulas and performs well in high dimensional cases. This package is not easily replaced in many other languages. The Julia language, on the other hand, has a far richer set of differential equation solvers than R or MATLAB^®^ and comes with very efficient forward sensitivity analysis methods.

Each researcher must therefore make decisions that result in the best compromise for them. If a researcher is familiar with a given set of programming/scripting languages it is probably not reasonable to expect them to be able to collaborate with other groups in languages they do not know. For this reason, it is our firm opinion that the conversion of models between different formats is a very important task and it is equally important to use formats that people can pick-up easily. To make format conversion flexible intermediate files are a great benefit which leads to a modular approach with possible validation between modules.

We also have to consider the FAIR data principles - the findability, accessibility, interoperability and reusability of data and associated infrastructure (Wilkinson et al. 2016). Here, we would like to address the interoperability principle by having developed a workflow for building biochemical pathway models using existing tools and custom-made, short, freely available scripts for the storage and refinement of models in all phases of development, ensuring interchangeability with other formats and toolkits at every step in the pipeline using standardized intermediate files (Fig. 1 and Table 1). Fig. 1 illustrates the relationship between the tools we used in this workflow. We re-estimated parameters of a previously developed model. We created a use case for others to reuse and modify that will be used to demonstrate how the depicted parts operate. In addition, we made two other use cases available for testing other parts of the workflow, with more information in the supplement. More extensive testing was done with a model of the mitogen-activated protein kinase (MAPK) cascade provided by the FindSim workflow (Viswan et al. 2018) and results can be found in the supplementary materials.

**Table 1:** An overview of available software packages

| Tool | Language | Purpose | Input Format(s) | Output Format(s) | Manual Tuning |
| --- | --- | --- | --- | --- | --- |
| **Pre-existing tools** |  |  |  |  |  |
| MATLAB® Simbiology® | MATLAB® | To simulate biochemical cascades | m, sbproj, SBML | m, sbproj, SBML | yes |
| MATLAB® Optimization Toolbox™ | MATLAB® | Optimization | none | none | no |
| COPASI | GUI | To set up and run COPASI simulations | SBML, CPS, SED-ML, COMBINE | SBML, CPS, SED-ML, C, COMBINE Archives, XPPaut, Berkeley Madonna | yes |
| NEURON | python | To simulate neuron models with added biochemical cascades | mod | not specific | yes |
| STEPS | python | Stochastic simulation of reaction-diffusion on tetrahedral meshes | python script | not specific | yes |
| NFsim |  | Network-free and hybrid simulation of rule based models | BNGL | not specific | yes |
| BioNetGen |  | Environment for rule-based models setup and simulation | BNGL | not specific | yes |
| VFGEN | C++ | Converts ODE vector field files (vf) to many other languages/formats | vf | .m, .R, .py, .c, many others | no |
| **Custom-developed tools** |  |  |  |  |  |
| Diagnostics tool | MATLAB® | Runs the model, compares it to the provided data and calculates scores | Sbtab as Excel | mat + figures | not if Sbtab is correctly formatted |
| Parameter estimation | MATLAB® | Parameter estimation using MATLAB® optimization tools | Sbtab as Excel | mat + figures | not if Sbtab is correctly formatted |
| GSA analysis | MATLAB® | GSA analysis using Sobol-Saltelli method implemented in MATLAB® | Sbtab as Excel | mat + figures | not if Sbtab is correctly formatted |
| Simulation Setup Webapp | python | Setup and run STEPS, BioNetGen and NFsim simulations | SBML, BNGL | BNGL, pySB | yes, removal of functional reaction rates and applying stimuli |
| get_thermodynamic_constraints.m | GNU Octave | Checks for thermodynamic constraints between parameters | m | Terminal text | no |
| **Conversion Tools** |  |  |  |  |  |
| Sbtab to SBML+m+tsv | MATLAB® | part of our MATLAB® toolchain | Sbtab as Excel | SBML, m, sbproj, tsv | no |
| sbtabs_to_vfgen.R | R | Convert Sbtab files into VFGENs vf format, but also produce SBML via libSBML | Sbtab as TSV or ODS | vf, SBML, mod | depends on the model and intended purpose |

**Fig. 1:**
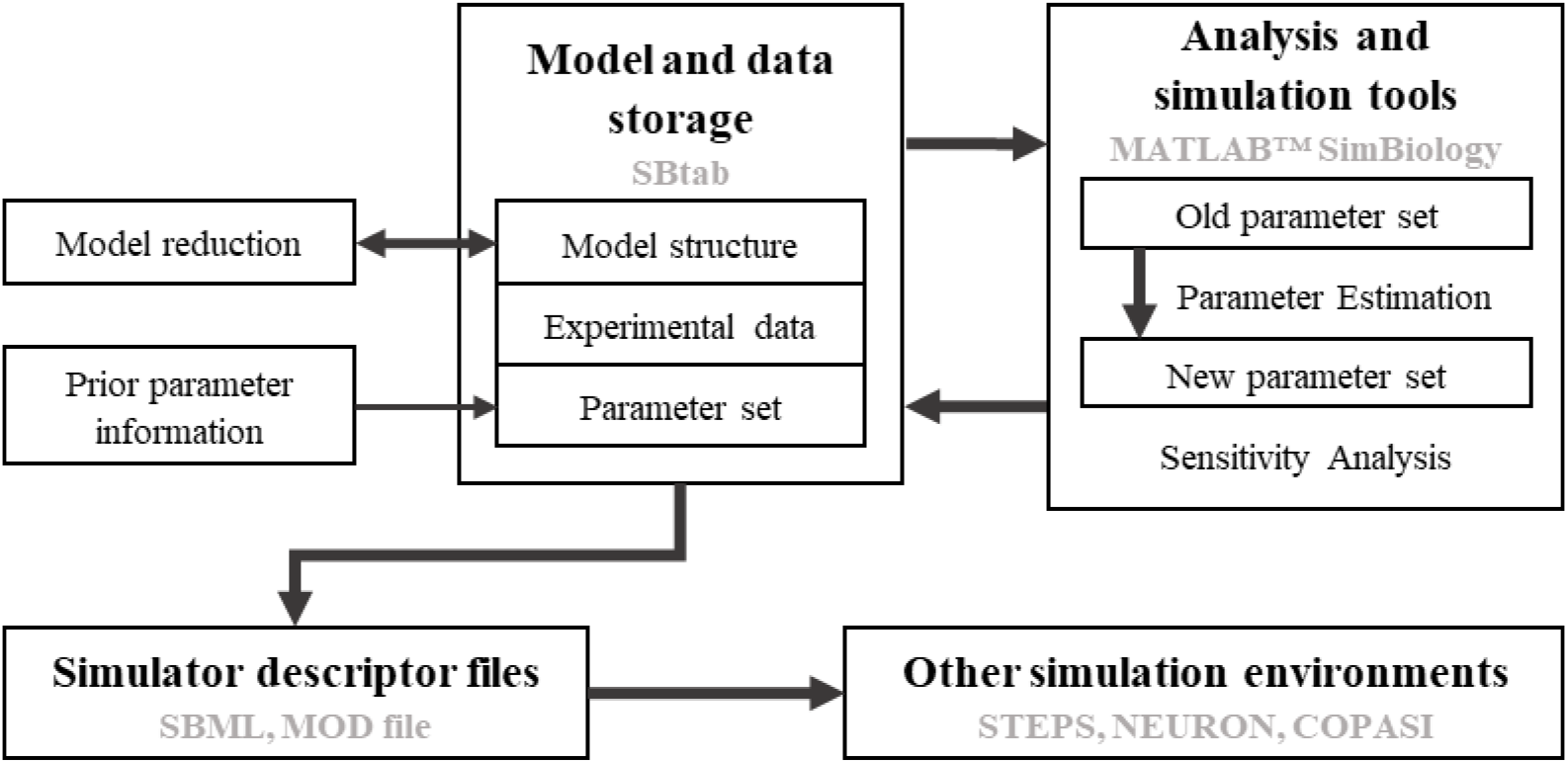
Simplified scheme of the workflow. Thick arrows indicate steps for which we have developed automated tools. Text in bold refers to the generic components of the workflow and text in grey refers to existing software and data formats used in the current version of the suggested workflow

## The workflow

While the standard model storage format in systems biology is SBML, it has some drawbacks: it does not lend itself to manual editing, the math in an SBML file is difficult to read and write manually, xml parsing is a difficult task that cannot be undertaken lightly by the novice programmer, species entries do not have a concentration unit attribute, time is handled very differently than any other model variable, etc. None of these issues are an error of course, but they are inconvenient for the inexperienced user. Therefore, this workflow is centered around building an easy-to-use infrastructure with models and data expressed in a spreadsheet-based storage format called SBtab (Lubitz et al. 2016). We chose SBtab as the primary modeling source file because it is human-readable and writable, it can contain both the model and the data, and because it is easy to write parsing scripts for it, such as a converter from SBtab to SBML using the libSBML^3^ interface in R. This ease of convertibility is used in the second focus of the workflow, convertibility between SBtab and other common formats and simulation software, since in systems biology and in any other computational sciences, the lack of compatibility between different tools and formats can often pose problems. A partially working conversion tool between SBtab and SBML had already been developed by the SBtab team. However, it can currently only read one table at a time and does not produce any functional SBML files with our model example. To combat these shortcomings, we wrote scripts to convert the SBtab into SBML, either using the R language or MATLAB^®^, and validating it successfully in COPASI, STEPS and NEURON.

The bulk of our workflow is available as MATLAB^®^ code, particularly the parameter estimation tools and functions for global sensitivity analysis. Sensitivity analysis can be used to determine the importance of different parameters in regulating different outputs. Local sensitivity analysis is based on partial derivatives and investigates the behavior of the output when parameters are perturbed in close vicinity to a specific point in parameter space. Tools for local SA are already included in the MATLAB^®^ suite. Global sensitivity analysis, on the other hand, is based on statistical approaches and has a much broader range. Global sensitivity analysis is more relevant for models that have a large uncertainty in their parameter estimates which is common for systems biology models where many of the parameters have not been precisely measured and the data are sparse.

The standard approach within biochemical modeling is to use deterministic simulations and ordinary differential equations (ODE) that follow the law of mass action as it is computationally efficient and provides accurate (compared to averaged stochastic simulations) results for sufficiently large well-mixed biological systems. However, this approach has several restrictions in the case of neuronal biochemical cascades. First, such cascades are always subject to stochastic noise, which can be especially relevant in a compartment as small as a dendritic spine where the copy number of key molecules are small enough that the effect of randomness becomes significant (Bhalla 2004). For precise simulation of stochasticity in reaction networks several stochastic solvers are available, e.g. Gillespie's Stochastic Simulation Algorithm (SSA) (Gillespie 1976) and explicit and implicit tau-leaping algorithms (Gillespie 2001). Second, the number of possible states of many biochemical cascades grow exponentially with the number of simulated molecule types, such that it becomes difficult to represent all these states in the model. In this case, for efficient simulation the reactions in the model could be represented and simulated in a network free form using rule-based modeling approaches (Chylek et al. 2015). Third, many biochemical networks are spatially distributed, this requires simulation of molecule diffusion (Hepburn et al. 2012). To tackle these problems, we developed the subcellular simulation setup application, a web-based software component for model development. It allows the extension and validation of deterministic chemical reaction network-based models by simulating them with stochastic solvers for reaction-diffusion systems (STEPS, Hepburn et al. 2012) and network free solvers (NFsim, Sneddon et al. 2011).

Although the workflow is applicable to any biochemical pathway model our emphasis is on modeling biochemical signaling in neurons. Therefore, the last challenge we want to address is an important concept in the interoperability domain of computational neuroscience called multiscale modeling which concerns the integration of subcellular models into electrical models of single cells or in neuronal microcircuits. This can be achieved either by run-time interoperability between two simulators of different systems or by expanding the capabilities of a single simulation platform as has been done with the NEURON software (McDougal et al. 2013). With this purpose in mind, we have written a conversion function from SBtab to the MOD format which is used by NEURON. As such, the inputs and the outputs of a biochemical cascade can be linked to any of the biophysiological measures of the electrical neuron model.

## Use case

As a primary use case to illustrate the workflow we have chosen a previously developed pathway model of the emergence of eligibility trace observed in reinforcement learning in striatal direct pathway medium spiny neurons (MSN) that carry the D1 receptor (Nair et al. 2016). Additional use cases are considered in the supplementary materials.

In this model, a synapse that receives excitatory input which leads to an increase in calcium concentration is potentiated only when the signal is followed by a reinforcing dopamine input. Fig. 2A represents a simplified model scheme illustrating these two signaling cascades, one starts with calcium as the input and the other one with dopamine. In simulation experiments the inputs are represented as a calcium train and a dopamine transient (Fig. 3A). Calcium input refers to a burst of 10 spikes at 10 Hz reaching 5 μM. Dopamine input is represented by a single transient of 1.5 μM. The first cascade (species in blue) features the calcium-dependent activation of Ca^2+^/calmodulin-dependent protein kinase II (CaMKII) and the subsequent phosphorylation of a generic CaMKII substrate which serves as a proxy for long term potentiation (LTP) and is the main output of the model. The second cascade (species in red) represents a G-protein dependent cascade following the dopamine input and resulting in the phosphorylation of the striatal dopamine- and cAMP-regulated phosphoprotein, 32 kDa (DARPP-32) that turns into an inhibitor of protein phosphatase 1 (PP1) which can dephosphorylate both CaMKII and its substrate. The phosphorylation of the substrate is maximal when two constraints are met. First, the time window between the calcium and dopamine inputs has to be short, corresponding to the input-interval constraint which is mediated by DARPP-32 via PP1 inhibition. Second, intracellular calcium elevation has to be followed by the dopamine input, corresponding to the input-order constraint that is mediated by another phosphoprotein, the cyclic AMP-regulated phosphoprotein, 21 kDa (ARPP-21), thanks to its ability to sequester calcium/calmodulin if dopamine arrives first (Fig. 3D).

**Fig. 2:**
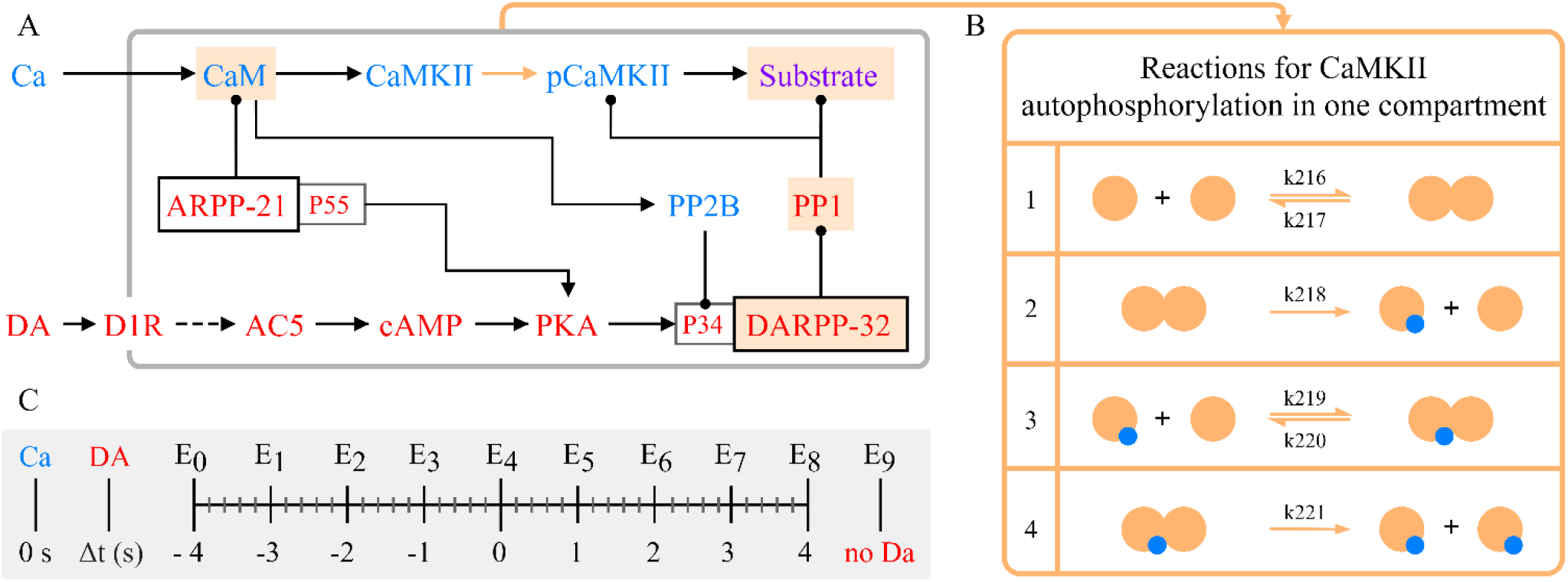
(A) Simplified schematics of the use case model with relevant second messengers with calcium and dopamine as inputs and phosphorylation of a generic CaMKII substrate (purple; top right) as the output. Species of the calcium cascade are blue, and species of the dopamine cascade are red. Lines ending in arrows represent activation and lines ending in circles represent inhibition. Time courses of the species with a beige background are later used in parameter estimation. Readjusted from Nair et al. (2016). (B) Schematics of the bimolecular reactions used for CaMKII autophosphorylation with yellow circles depicting fully activated CaMKII bound to calmodulin and calcium, and blue circles depicting phosphate groups. Six newly introduced parameters are shown on the reaction arrows with their ID’s in the updated model. (C) Timing (in seconds) of the dopamine input (Δt= {−4,−3,−2,−1,0,1,2,3,4} corresponding to E0-E8) relative to the calcium input (zero), and a single experiment without a dopamine input (E9)

**Fig. 3:**
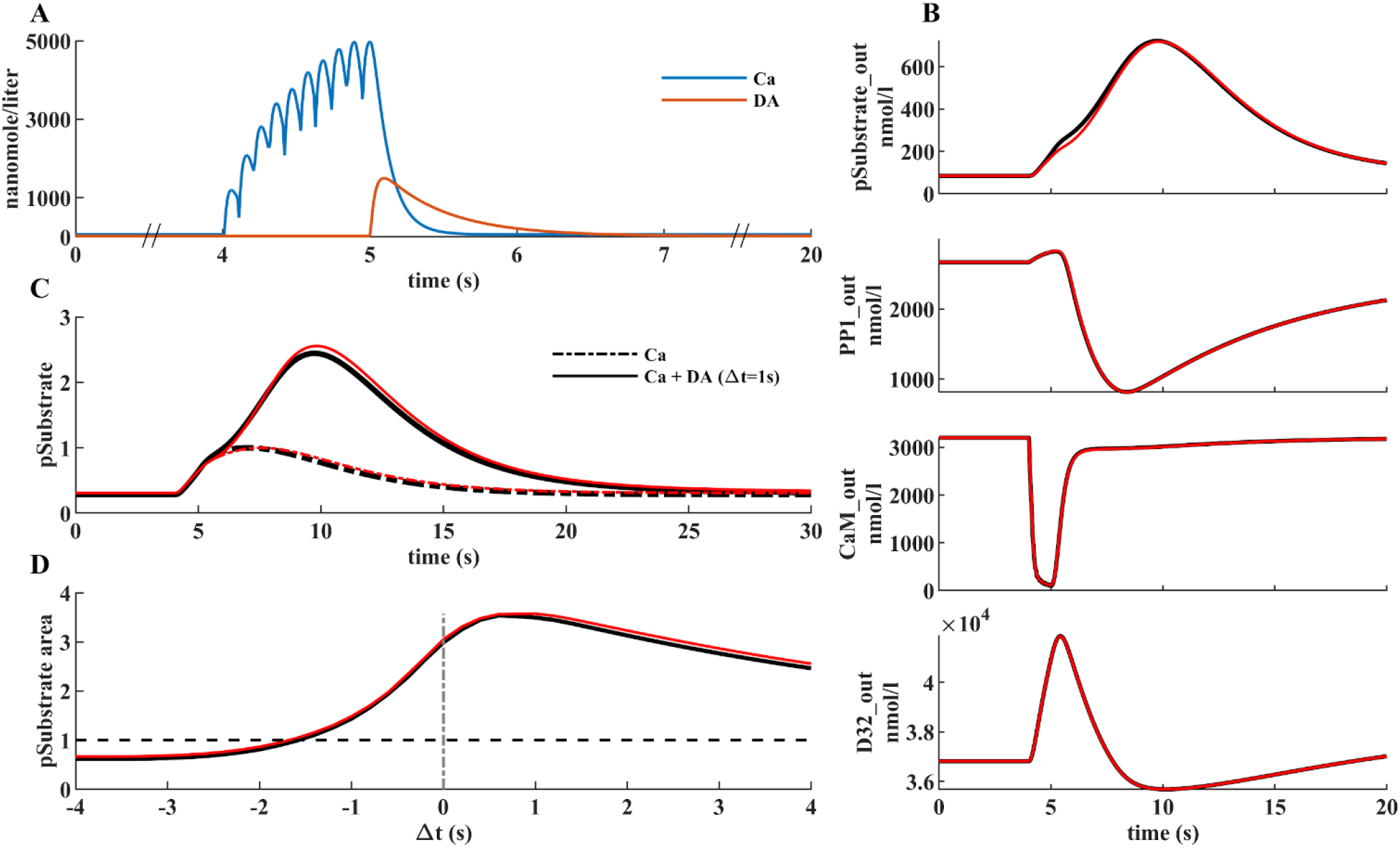
(A) Illustration of the model inputs. Calcium burst (blue) at 4 s used in all simulations and a dopamine transient (orange) applied at different timings in eight experiments and one without it. (B) Four species used in parameter estimation corresponding to the input combination in A. Black traces represent the data produced by simulating the original model, red traces represent fits with the best new parameter sets in the updated model. (C, D) Comparison of model performance with substrate phosphorylation as the main model readout. (C) Normalized time series of substrate phosphorylation, the main readout, with calcium as an only input or dopamine following it after 1 s. (D) Normalized area under the curve of substrate phosphorylation with different calcium and dopamine input intervals

In the originally published model, CaMKII is autophosphorylated in two compartments, both the cytosol and the post synaptic density (PSD), with a custom-written MATLAB^®^ rate function that was calculated based on the probability of two neighboring subunits being fully activated as described in Li et al. (2012). To make it possible to run the model in different software we replace the rate equation of autophosphorylation with a similar set of reactions in both compartments so that the model would only contain bimolecular reactions. The reactions represent a simplified version of the autophosphorylation reactions in Pepke et al. (2010), where in our case only the fully activated CaMKII can be phosphorylated. The same set of reactions is used in both compartments and the schematics is available in Fig. 2B along with the required six new parameters. We used our parameter estimation script to find parameter values and bounds that preserved the qualitative behavior of the model. In this primary use-case, we used simulated data from the original model with different timings of the dopamine input relative to the calcium input (Fig. 2C) to obtain a comprehensive picture of its behavior which we want the updated model to reproduce.

## SBtab

As described above we have chosen SBtab as the format at the root of our workflow for the storage of model and data. In this section we illustrate how we use this format, for more information, documentation and examples are available from the SBtab authors^4^. This format allows the storage of biochemical models and associated data in a single file and provides a set of syntax rules and conventions to structure data in a tabulated form making it easy to write, modify and share. To ensure interoperability, SBtab provides an online tool to convert the models into the SBML format^5^. SBtab is suitable for storing data that comes in spreadsheet or table formats, e.g. concentration time series or dose response curves, but it is likely that *any* data format that can be reasonably stored as a table will work well in SBtab. The SBtab file is intended to be updated manually during the process of model building. Additional instructions on how to make SBtab files work well within our toolchain can be found in the Subcellular Workflow documentation^6^. Some of the columns and sheets that we use should be considered as extensions to the format and are discussed in the documentation. SBtab is easy to parse so adjustments to parsers can be made quickly.

The SBtab file should include separate sheets for compartments, compounds, reactions, assignment expressions, parameters, inputs, outputs, and experiments (as well as data tables). We illustrate the functionalities of SBtab here with one use case. The use case model has 99 compounds, 138 reactions and 227 parameters. An example of the SBtab reaction table can be found in Table 2.

**Table 2:** An example of the tabulated representation of model reactions in the SBtab format. The kinetic law uses the parameter names from the parameter table and species names from the compound table.

| !ID | !Name | !KineticLaw | !IsReversible | !Location | !ReactionFormula |
| --- | --- | --- | --- | --- | --- |
| R0 | ReactionFlux0 | kf_R0*GaolfGTP | FALSE | Spine | GaolfGTP <=> GaolfGDP |
| R1 | ReactionFlux1 | kf_R1*D1R_Golf_DA | FALSE | Spine | D1R_Golf_DA<=>Gbgolf+D1R_DA+GaolfGTP |
| R2 | ReactionFlux2 | kf_R2*D1R_Golf*DA-kr_R2*D1R_Golf_DA | TRUE | Spine | D1R_Golf+DA<=>D1R_Golf_DA |
| R3 | ReactionFlux3 | kf_R3*D1R*DA-kr_R3*D1R_DA | TRUE | Spine | D1R + DA <=> D1R_DA |
| R4 | ReactionFlux4 | kf_R4*AC5*GaolfGTP-kr_R4*AC5_GaolfGTP | TRUE | Spine | AC5 + GaolfGTP <=> AC5_GaolfGTP |
| R5 | ReactionFlux5 | kf_R5*CaM*Ca-kr_R5*CaM_Ca2 | TRUE | Spine | CaM + Ca <=> CaM_Ca2 |
| R6 | ReactionFlux6 | kf_R6*PP2B*CaM-kr_R6*PP2B_CaM | TRUE | Spine | PP2B + CaM <=> PP2B_CaM |

One of our goals with this study was to reproduce the original model behavior after replacing a single module inside the model to convert it to bimolecular reactions only. The data we used therefore represents the simulated time series (20 s) of the concentrations of four selected species in response to different input combinations using the original model. Each individual data sheet (named E0-E9, Fig. 2C) in SBtab represents the outputs of one experiment. Another sheet called Experiments allows to define the input parameters differently for each experimental setup. By setting the initial concentrations of the unused species to zero the data could be mapped to a specific sub-module of the model (the remaining species). In this case the initial conditions are the same for all experiments. The Experiments table can also support annotations relevant to each dataset. We used nine different timings (corresponding to E0-E8) between the calcium and the dopamine signal starting with a dopamine signal preceding calcium by four seconds and finishing with dopamine following calcium after four seconds as this corresponds to the time frame originally used in model development (Δt={−4,−3,−2−1,0,1,2,3,4}). Additionally, we used simulations with calcium as the only input (E9) (Fig. 2). The time series of the input species are in a separate sheet following each experiment sheet. An example of how experimental data is stored can be found in Table 3.

**Table 3:** An example of the tabulated representation of experimental time series data in the SBtab format. The table contains the data for experiment E0. Columns titled Y0-Y3 refer to each of the “measured” output species followed by a standard deviation column. The time series in this data each contain 2001 data points.

| !TimePoint | !Time | >Y0 | SD_Y0 | >Y1 | SD_Y1 | >Y2 | SD_Y2 | >Y3 | SD_Y3 |
| --- | --- | --- | --- | --- | --- | --- | --- | --- | --- |
| E0T0 | 0 | 84.47891 | 1 | 2672.089 | 1 | 3200.526 | 1 | 36817.52 | 1 |
| E0T1 | 0.01 | 84.47891 | 1 | 2672.089 | 1 | 3200.526 | 1 | 36817.52 | 1 |
| E0T2 | 0.02 | 84.47891 | 1 | 2672.089 | 1 | 3200.526 | 1 | 36817.52 | 1 |
| E0T3 | 0.03 | 84.47891 | 1 | 2672.089 | 1 | 3200.526 | 1 | 36817.52 | 1 |
| ... |  |  |  |  |  |  |  |  |  |
| E0T2000 | 20 | 97.79736 | 1 | 2359.101 | 1 | 3192.558 | 1 | 37475.51 | 1 |

## Model pre-processing tools

Model building entails frequent changes to the model structure by adding new species, reactions, and parameters. This can result in the emergence or disappearance of Wegscheider cyclicity conditions that refer to the relationships between reaction rate coefficients arising from conditions of thermodynamic equilibrium (Wegscheider 1901; Vlad and Ross 2004). Identified thermodynamic constraints show parameter dependencies that follow from physical laws and can reduce the number of independent parameters. These conditions are frequently difficult to determine by human inspection, especially for large systems. Similarly, identifying conserved moieties, like conserved total concentration of a protein, allows the reduction of the ODE model size, which leads to increased performance. In order to address these model pre-processing needs our toolkit includes scripts in MATLAB^®^/GNU Octave that use the stoichiometric matrix of the reaction network as an input to determine the thermodynamic constraints as described in Vlad and Ross (2004), and conservation laws. These diagnostic tools output any identified constraints that, if needed, are to be implemented manually before the parameter estimation step. It should be noted that such constraints need to be re-examined after each addition of new reactions as the structure of the model might change and make previously true constraints invalid. This is true for all major changes to the model.

## MATLAB^®^ tools

The bulk of our workflow is developed in MATLAB^®^ as it provides an easy-to-use biochemical modeling application with a graphical user interface called SimBiology^®^ along with a wide range of toolboxes for mathematical analysis. The workflow is divided into import and analysis scripts. We have written software for three types of analysis: diagnostics tools, used to run the model, inspect the data and model representation of the data, parameter estimation, and global sensitivity analysis. To ensure an easy and user-friendly usage, all operations are controlled by a single settings file where all specification options needing user input are represented as modifiable variables. An example settings file of the use case model along with instructive comments can be found in the GitHub repository^7^. Only this settings file and the SBtab file are needed as input from the user to run all our MATLAB^®^ scripts. After running an analysis, the results are stored inside the model folder and relevant plots are generated. This process is entirely automatic, but the user can always explore and retrieve more data from the created files. For example, after running the parameter estimation analysis, a plot is generated with the original parameter set, the prior bounds, and optimized parameters, but the procedure for retrieving the optimized parameters and using them to create a new SBtab or a new settings file for subsequent runs is not yet automated. To get a full sense of what is plotted and what kinds of data are created but not immediately shown please see our GitHub repository documentation^8^.

## Import from SBtab to MATLAB^®^

The import scripts we have written generate all the files that are needed for proper running of our MATLAB^®^analysis, saving them in subfolders of the main model folder. Among these files there is a version of the model without any inputs in the MATLAB^®^ (.mat) and SimBiology^®^ (.sbproj) format, and several versions of the model corresponding to each experiment defined in the SBtab, with the model ready to be equilibrated and with the model ready to be simulated after equilibration in the MATLAB^®^ (.mat) format. The versions ready to be simulated include all the inputs and outputs to be measured as specified in the SBtab but not the starting values, as the equilibrations are only performed at run time. Additionally, while not needed for the rest of our MATLAB^®^ workflow, these import scripts also generate .tsv files corresponding to the SBtab (useful for tracking changes in GitHub), an SBML file using MATLAB^®^ built-in functions (level 2 version 4 encoding as of MATLAB^®^ 2020b) useful for exporting the model into other simulators (note that we have another converter from SBtab into SBML that runs in R instead of MATLAB^®^).

## Parameter estimation

MATLAB^®^ offers a wide range of tools for function optimization. We have developed scripts that transform our parameter estimation problem into an objective function that can be optimized by various MATLAB^®^ built-in optimizer functions. At the time of writing, these include “fmincon”, simulated annealing (“simulannealbnd”), pattern search (“patternsearch”), genetic algorithm (“ga”), particle swarm (“particleswarm”), and surrogate optimization algorithms (“surrogateopt”), for which MATLAB^®^ provides thorough documentation. The code is built with flexibility in mind, so introduction of other MATLAB^®^ built-in or custom optimization algorithms should be straightforward. The optimizers used are the ones chosen in the settings file, our code supports the use and comparison of multiple optimizers at the same time, and multiple uses of the same optimization algorithm are also supported. This is particularly useful for using optimizers that are inherently single-threaded, e.g. the simulated annealing algorithm, since multiple simulated annealing optimizations can be performed in multiple computing cores. After performing the optimization, a file containing the results is outputted then from here the optimized outputs can be taken and used to manually update the SBtab or SimBiology^®^ model. One of the equations used to calculate the score for how well the model outputs fit the experimental data can be found below (Equation 1). We have incorporated a few other ways of calculating the score, and custom scoring methods could also be added depending on the need (see documentation).

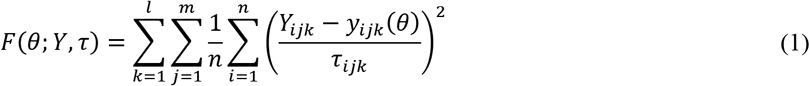

Here, *Y* represents the data that is going to be used to constrain the model, sourced either from experiments or previous models, and *y* is the outputs of the model mapped to the data resulting of the simulation of the model under parameterization *θ*. The allowed mismatch *τ* between the two simulation results is analogous to the standard deviation of a Gaussian noise model in data fitting. The resulting *F* is the objective function for optimization. The error is summed over *n*, the number of points in each experimental output, *m*, the number of experimental outputs in an experiment (which is four in our use case, see Fig. 3B), and *l*, the number of experiments (E0-E9 in our use case) (see Fig. 2C).

Parameter estimation is generally based on experimental data. In this use case, however, we used simulated experimental data of the concentrations of several species using the original version of the model in SimBiology^®^ (two additional use cases are provided in the Supplementary Materials using actual experimental data). After modifying the model, we minimized the difference between the old behavior and the updated model’s response through optimization. The simulation results from the old model can be considered as analogous to experimental data in a normal parameter estimation setting. Here, we merely aim to make an updated model agree with its earlier iteration, which itself was adjusted based on experimental data. When changing a module in a model it is crucial to protect the unchanged parts, which is why we performed parameter estimation using the key species that intersect the calcium and dopamine cascades, namely PP1, calmodulin and DARPP-32 (Fig. 2 and Fig. 3B). In this use case, the Particle Swarm Algorithm was chosen to perform the optimization, but all algorithms were capable of reasonable optimizations. The parameters obtained were then used to generate all the figures where optimized parameters are referred. The choice of total amount of reactions, used to replace the original function that represented the CAMKII phosphorylation, were constrained by the optimization. We considered the outputs that we were measuring (Fig. 3B) and added reactions until the addition of more did not meaningfully improve the fits.

## Validation in MATLAB^®^

We developed diagnostics scripts that can be used to reproduce the various experiments defined in the SBtab. These scripts generate plots of the experimental inputs to the model (adapted for Fig. 3A), the provided data and the outputs measured from model simulation (adapted for Fig. 3B) given some choice of parameters, and plots of the scores calculated for the differences between the various experimental outputs and simulated model outputs. We used these tools to confirm that our parameter estimation resulted in a good fit for most of the species and the updated model was able to closely reproduce the results seen with the original model (Fig. 3C). In our repository we provide the updated model in SBtab (.xlxs and .tsv), SBML (.xml) and MATLAB^®^ SimBiology^®^ (.sbproj and.mat). In addition to the general-purpose tools, we also wrote a use case-specific script, which uses data from the original model and reproduces the time-dependency of the substrate phosphorylation given different delays of the start of calcium and dopamine stimuli, using the optimized model (Fig. 3C–D).

## Global sensitivity analysis

In many cases parameter estimation of biochemical pathway models does not result in one unique value for a parameter. Structural and practical unidentifiability (Raue et al. 2009) results in a large set of parameter values that all correspond to solutions with a good fit to the data, i.e., there is a large uncertainty in the parameter estimates (Eriksson et al. 2019). When this is the case, *local* sensitivity analysis is not so informative, since this can be different depending on which point in parameter space it is performed at. A global sensitivity analysis (GSA), on the other hand, covers a larger range of the parameter space. Several methods for GSA exist (Zi 2011) but we have focused on a method by Sobol and Saltelli (Sobol 2001; Saltelli 2002; Saltelli 2004) as implemented by Halnes et al. (2009) which is based on the decomposition of variances (Saltelli 2004). Single parameters or subsets of parameters that have a large effect on the variance of the output get a high sensitivity score in this method. Intuitively, this method can be understood as varying all parameters but one (or a small subset) at the same time within a multivariate distribution to determine what effect this has on the output variance. If there is a large reduction in the variance, the parameter that was kept fixed is important for this output (Saltelli 2004).

Let the vector *Θ* denote the parameters of the model, and *y=f(Θ)* be a scalar output from the model. In the sensitivity analysis *Θ* are stochastic variables, sampled from a multivariate distribution, whose variation gives a corresponding uncertainty of the output, quantified by the variance *V(Y)*. Note that in the setting and interpretations described here, the different *Θ_i_* are assumed to be independent from each other (for cases with dependent *Θ_i_* see e.g. (Saltelli 2004) and (Eriksson et al. 2019)). We consider two types of sensitivity indices: the *first order effects S_i_* and the *total order effects S_Ti_*. The first order effects describe how the uncertainty in the output depends on the parameter *Θ_i_,* i.e., how much of the variance of the output can be explained by the parameter *Θ_i_* alone. As an example, *S_i_*=0.1 means that 10% of the output variance can be explained by *Θ_i_* alone. The total order effects give an indication on the interactive effect the parameter *Θ_i_* has with the rest of the parameters on the output. Parameters are said to interact when their effect on the output cannot be expressed as a sum of their single effects on the output.

The first order sensitivity index of the parameter *Θ_i_* is defined as

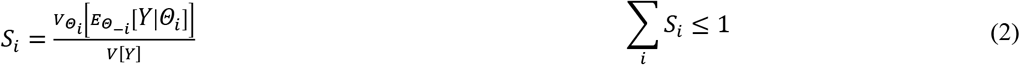

where *Θ_−i_* corresponds to all elements of *Θ* except *Θ_i_*. The total order sensitivity index of the parameter *Θ_i_* is defined as^9^

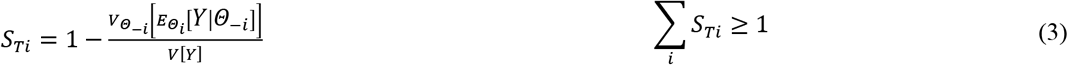

If there is a large difference between S*_i_* and S_Ti_, this is an indication that this parameter takes part in interactions. For a detailed description see chapter 5 of Saltelli (2004).

The optimization described earlier takes place on log transformed space (log_10_(*Θ*)). For the sensitivity analysis we perform the sampling on a lognormal distribution, meaning that log_10_(*Θ*)~ *N(μ, σ).* Below we use *μ=*log_10_(*Θ\**) and *σ*=0.1, where *Θ\** correspond to the optimal values received from the optimization^10^. We illustrate this method using only the six parameters corresponding to the model module that has been replaced (Fig. 2) and the results can be seen in Fig. 4 Only four of the parameters (k216-k219/k222-k225; Fig. 4) seem to be important for the output within the investigated parameter region. In experiments E0 and E1 these four parameters are shown to have approximately equal importance, whereas in experiments E3-E9 the parameter k219/k225 has the largest influence. Also, there seem to be some interactive effects between the parameters since S_Ti_ is larger than S_i_, especially for the first four experiments (Fig. 4A and B).

**Fig. 4:**
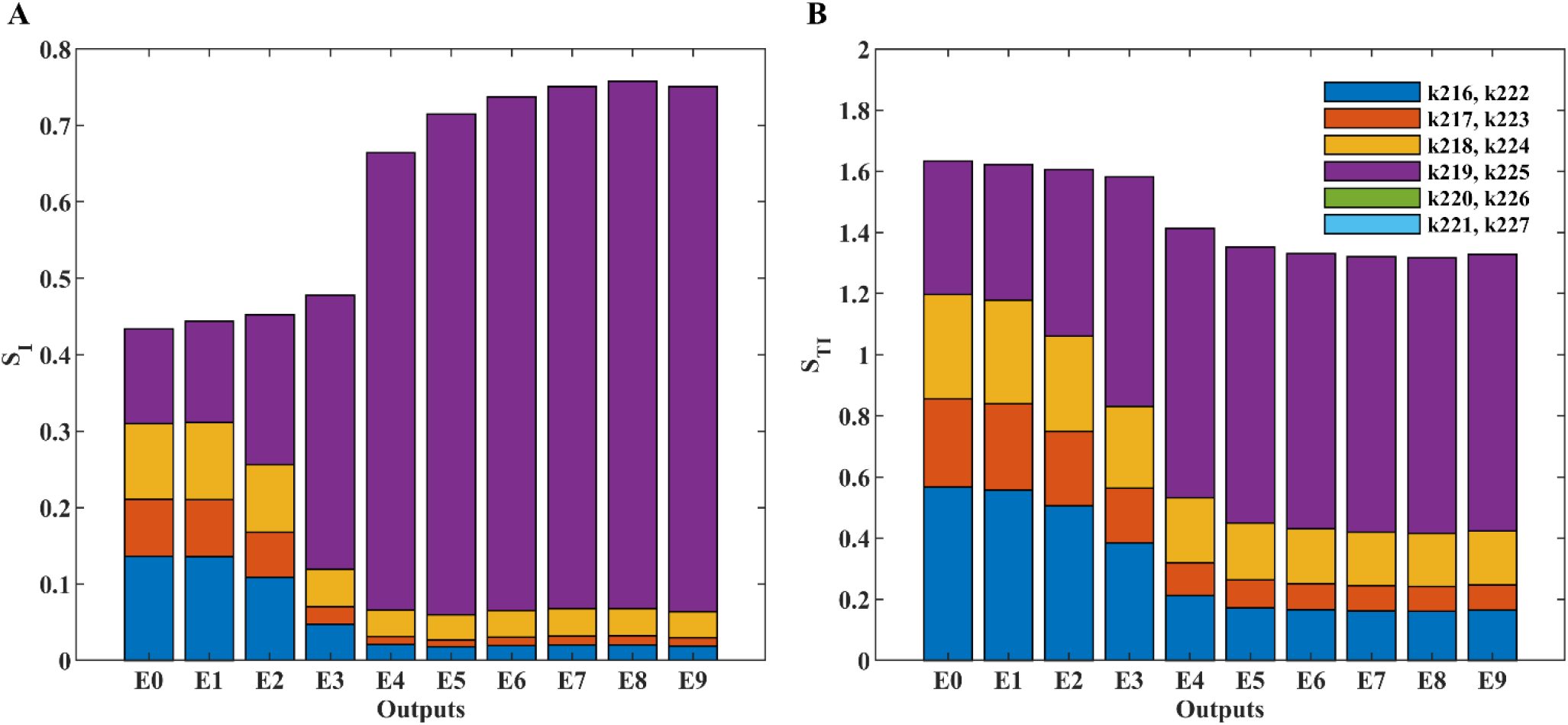
Stacked bar graphs of the sensitivity indices. The first order, Si, and total order, STi, sensitivities indices of the six new parameters (k216-k221 and k222-k227; see Fig. 2) for all ten experiments (E0-E9) are shown (panel A and B, respectively). The sensitivities indices are defined in the main text and were calculated based on the scores used in the optimization for each experiment respectively. The parameters, Θ, were sampled independently from a multivariate lognormal distribution with log10(Θ)~ N(μ, σ), using μ =log10(Θ*) and σ=0.1, where Θ* correspond to the optimal values received from the optimization. A sample size of N=10000 was used (corresponding to 80000 reshuffled samples used in the calculations (Saltelli 2004)). The analysis took 170 minutes on 50 compute cores (Intel Xeon E5-2650 v3). The sample size was chosen big enough to make the differences in the sensitivity scores stemming from different seeds11 small for the purposes of our conclusions.

## Compatibility and validation with other simulation environments

### Conversion to SBML and simulations in COPASI

COPASI is one of the more commonly used modeling environments in systems biology and it can read SBML files (Hoops et al. 2006). Our first validation step is to use the SBML model retrieved from the SBtab model in COPASI. The online conversion tool from SBtab to SBML did not work for our specific use case; we wrote a new conversion function that can be found in the GitHub repository^12^ of the paper. It interprets the biological model and converts it into plain ODEs in VFGEN’s custom format (.vf), the VFGEN file can then be used to create output in various languages^13^ (Weckesser 2008). The conversion script (written in R) converts the SBtab saved as a series of .tsv files or one .ods file into a VFGEN vector field file and as by-products also the SBML and a MOD file (see chapter *Conversion to a MOD file and simulations in NEURON*). To create an SBML model libsbml must be installed with R bindings. The SBML file can be imported directly into COPASI, although it might be necessary to remove the superfluous unit definitions manually.

Another way to convert the model into SBML is through a single MATLAB^®^ SimBiology^®^ function. The models, however, are created with long ID’s that carry no biological information (the IDs are similar to hexadecimal hashes) for all model components, the units are not properly recognized, and some units may just be incorrectly defined in the output. We, therefore, created a script in R that asks for default units for the model and replaces the ones in the SBML file. It fixes most issues (units, IDs, and the time variable in assignments) allowing the model to be properly imported in^14^ COPASI. To illustrate that the SBML-converted model imported into COPASI produces the same results as in SimBiology^®^, we used a simplified calcium input corresponding to one double exponential spike analogous to the dopamine transient and simulated the model with deterministic solvers in both COPASI (LSODA solver) and MATLAB^®^ SimBiology^®^ under similar conditions. Both simulation environments produced almost overlapping results (Fig. 5B-C) validating the converted model in the SBML format.

**Fig. 5:**
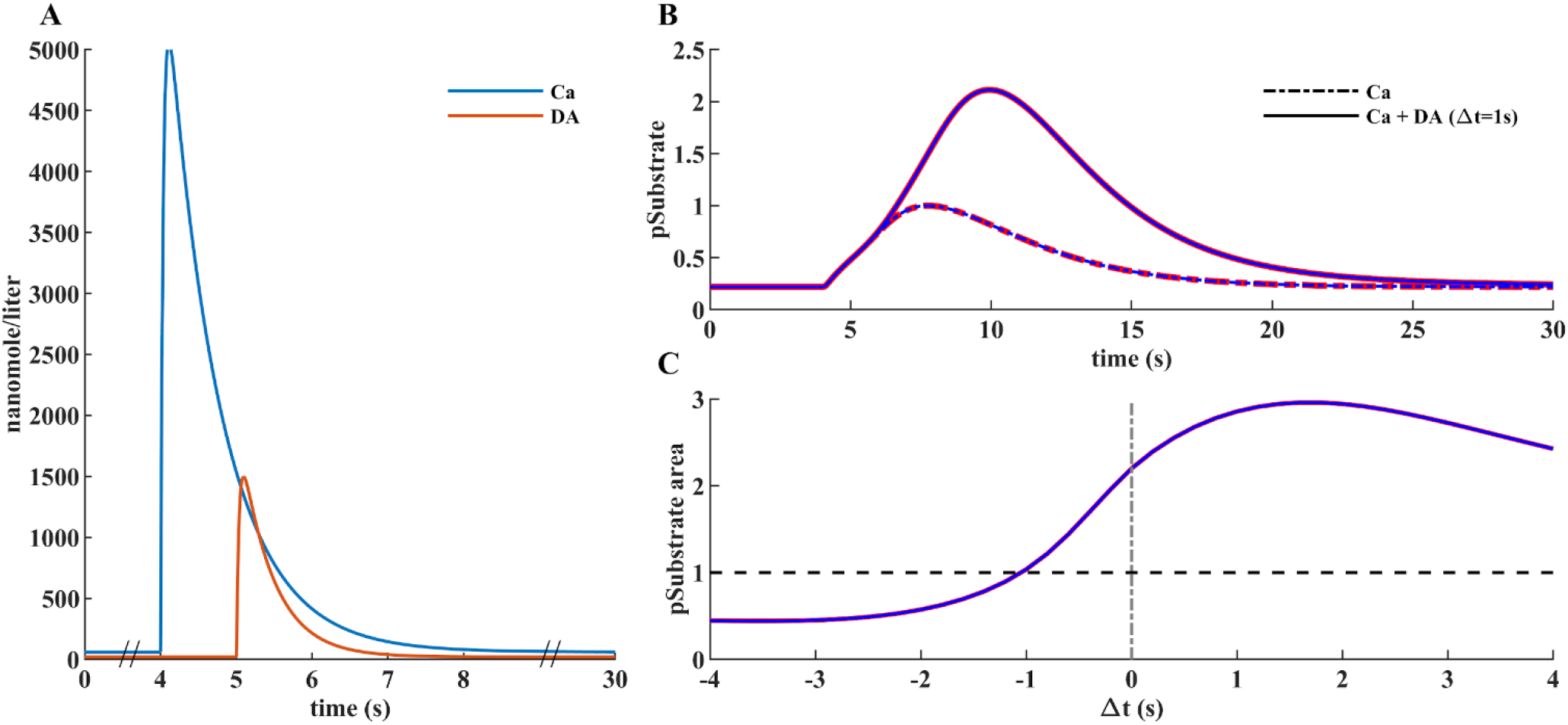
Simulations in identical conditions in both MATLAB^®^ SimBiology^®^ and COPASI yielded almost identical results. (A) Inputs used in both simulators. The calcium input is kept constant at 4 s for all simulations and dopamine input time is varied from time 0 to 8 s at every one second. The difference from the previous simulations is in the calcium input which, for the sake of simplicity, is represented by a double exponential spike. (B) and (C) show substrate phosphorylation curves analogous to Fig. 3, the red line represents results obtained in MATLAB^®^ and blue line results from simulations in COPASI.

## Simulations in STEPS

The web-based subcellular simulation setup application^15^ allows importing, combining and simulating models expressed in the BioNetGen language (BNGL; Harris et al. 2016). It supports the import of SBML (level 2 version 4) models and their transformation to rule-based BNGL form using Atomizer (Tapia and Faeder 2013). The BioNetGen file format was extended to provide diffusion parameters, links to tetrahedral meshes describing the geometry of model compartments, as well as the additional parameters for solvers and stimulation protocols required for spatially distributed models. The subcellular simulation setup application is integrated with the network free solver NFsim (Sneddon et al. 2011) and it supports simulations of spatially distributed systems using STEPS (Hepburn et al. 2012). STEPS provides spatial stochastic and deterministic solvers for simulations of reactions and diffusion on tetrahedral meshes. Furthermore, the subcellular simulation setup application provides a number of facilities for the visualization of models’ geometries and the results of simulations.

To demonstrate the compatibility of the subcellular simulation setup application with the workflow for model development described above, we imported the SBML version of the use case model to the setup application and simulated it with the STEPS TetOpSplit solver. We have used a simple two-compartmental spine model with a tetrahedral compartment corresponding to the spine compartment of the use case model. There is also a PSD compartment on one of the faces. The results of the model simulations with a STEPS solver were qualitatively similar to the results obtained with the deterministic model simulated in MATLAB^®^. Examples of simulated time courses for molecule concentrations as shown in Fig. 3 in comparison with corresponding MATLAB^®^ curves are shown in Fig. 6.

**Fig. 6:**
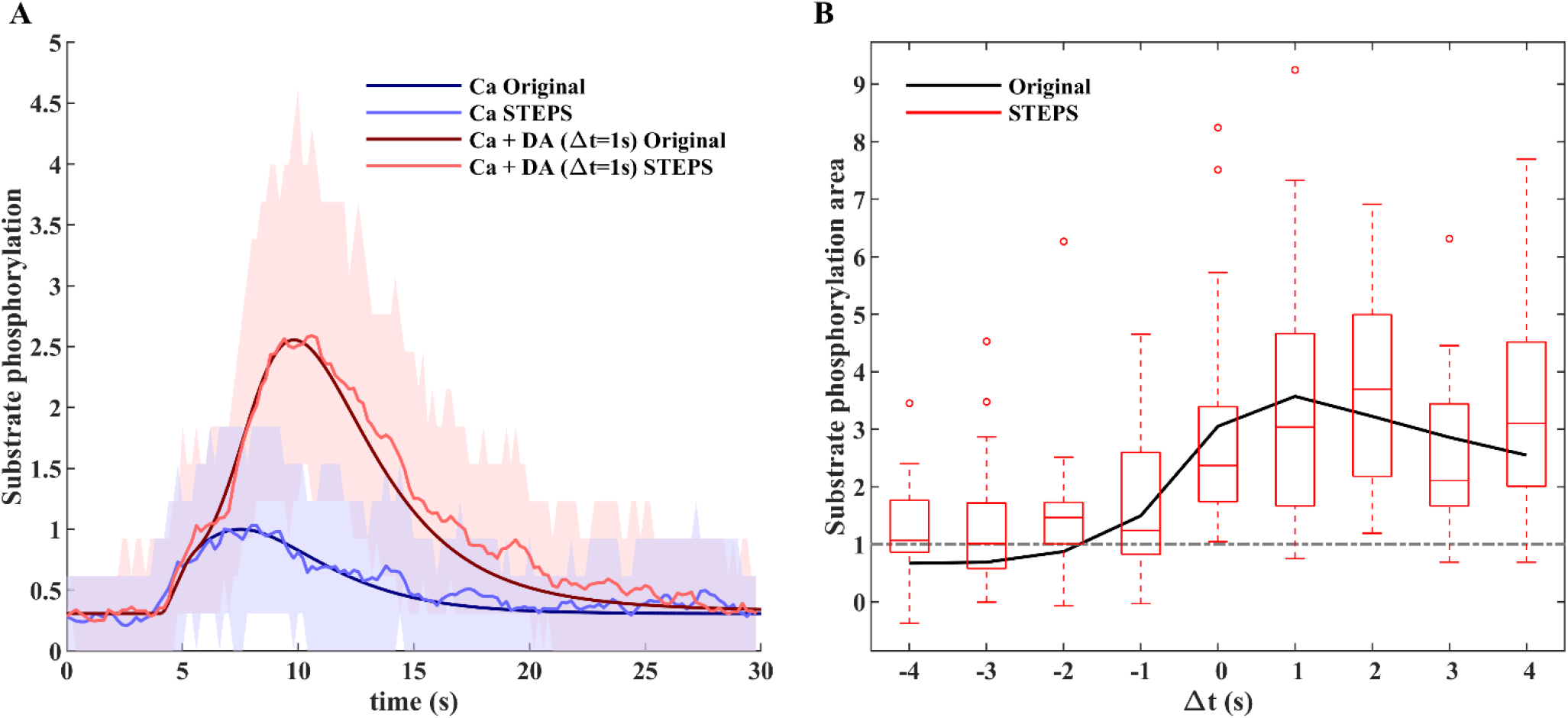
Validation of the model by stochastic STEPS simulation of substrate phosphorylation in a typical D1 MSN neuron spine. (A) Normalized time course of substrate phosphorylation in the original model in comparison with averaged (n=50) stochastic STEPS simulations (red - calcium and dopamine; blue - calcium only) for a typical size of D1 MSN neuron synaptic spine (V=0.02μm3). The same stimulation protocol as in Fig. 3 was used. Colored areas around averaged STEPS curves correspond to a range between 10% and 90% confidence intervals. (B) Normalized area under the curve of substrate phosphorylation with different calcium and dopamine input intervals simulated for the MATLAB^®^ version of the updated model (with MATLAB^®^ ode15s solver, black line) and averaged stochastic STEPS simulations (n=30) in the application version of the model. MATLAB^®^ statistical bar plots were added to the figure to characterize variability of synaptic plasticity between subsequent induction protocol applications to the same synaptic spine. Note that despite high variability of synaptic plasticity time courses averaged plasticity dynamics were in a good agreement with the ODE-based solution.

## Conversion to a MOD file and simulations in NEURON

As we suggested before, conversion between different modeling frameworks and formats facilitates collaboration. But conversion is made harder by the differences in the capabilities of different modeling packages. A model, such as the one above, could be useful for multiscale simulations investigating how neuronal network activity shapes synaptic plasticity. Many cellular level models are built and simulated in the NEURON environment which also supports simplified reaction-diffusion systems. It is useful to be able to integrate a subcellular level model into a cellular level model specified using NEURON. Models in NEURON are built by adding features with MOD files that are written in the NMODL programming language. A schematic illustration of our use case set up in NEURON is depicted in Fig. 7. Conversion from SBML to MOD is already possible via NeuroML (instructions can be found in Lindroos et al. 2018); it is not an automated or user-friendly approach. We wrote an R script, which reads an SBtab model^16^ and writes a MOD file, as well as VFGEN and an SBML^17^ file. The script can optionally perform analysis of conservation laws and output a model where some of the differential equations for the state variables are substituted by algebraic equations arising from the conservation laws, thereby reducing the number of differential equations to be solved. An example on how to use the SBtab to VFGEN/MOD/SBML converter in R can be found in the GitHub repository^7^. The subcellular model in SBtab form is not aware of its coupling to a larger model of the cell and the user must edit the resulting MOD file manually to use it within a larger scope and assign a role to this model component. This typically means assigning input to the model and using the output in some way. An example is given below with our use case. Another point to stress is that the script converts the time unit of the parameters to milliseconds, NEURON’s default unit for time, but does not change the concentration units. Thus, when coupling a biochemical cascade to other quantities in the neuronal model, care must be taken to rescale the coupled variables so that they match the units in the rest of the neuronal model. We also illustrate the with one of the inputs in the use case.

**Fig. 7:**
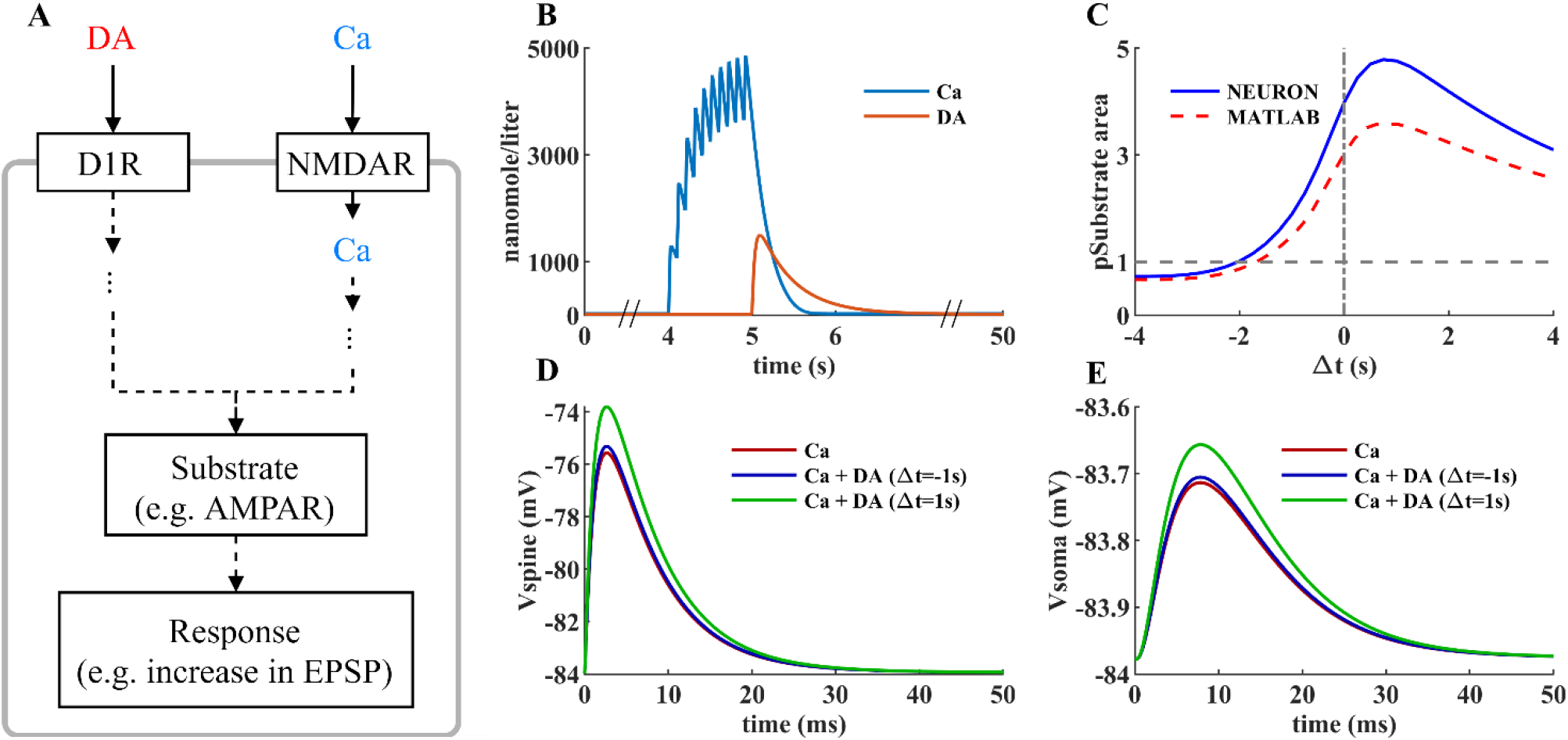
Inserting the biochemical signal transduction cascade into an electrical model in NEURON. (A) A schematic of the effects of the two inputs of the model, dopamine and calcium, on a generic substrate, which in this case is taken to represent the fraction of phosphorylated AMPA receptors with higher conductance levels. (B) Examples of the two inputs, calcium and dopamine. The large calcium signal is due to an NMDA spike evoked via clustered excitatory synapses on a dendritic branch. (C) The calcium input from NMDA receptors also qualitatively reproduces the timing dependence obtained with the simulated model in MATLAB (result from Fig. 3D). (D, E) Predicted EPSP following a single synaptic input in the relevant spine and in the soma. The readout of the substrate phosphorylation level was done at 7s after the start of the calcium input. The relative timings of dopamine and calcium indicated with arrows in (C) are used, and the results are compared to the experimental setting without the dopamine input.

We validate the biochemical cascade model and our conversion tools in NEURON by qualitatively reproducing the results obtained in MATLAB^®^ SimBiology^®^. Our goal is to show that the cascade model can be integrated into a single neuron model and bridge spatial and temporal scales of system behavior by linking the output of the cascade to changes in the synaptic properties and ultimately to the electrical behavior of the neuron model. Therefore, the biochemical model in the MOD format was incorporated into a single biophysically detailed and compartmentalized D1 MSN model from Lindroos et al. (2018). To integrate the MOD file into single cell models a few user-specific modifications have to be made to interface this model with the larger electrochemical system. First, the input is modified so that the calcium burst is represented by calcium influx from the cell’s calcium channels and adjusted so that the overall calcium level would be similar to the input used in MATLAB^®^ simulations. The calcium in the neuron model is expressed in millimolar units and when coupling it to the biochemical cascade we rescale it to nanomolar units (the units used in the biochemical cascade model). The dopamine transient is represented by an assignment expression (available in the Expression table of the SBtab) that creates its double exponential form (this can be used in other languages as well). When simulating the model with the same input timing combinations as in the MATLAB^®^ experiments, we were able to qualitatively reproduce the substrate phosphorylation curve that illustrates both the input-order and input-interval constraints seen in Fig. 3D. The differences can be explained by differences in the calcium input (minor differences can also arise due to the different numerical solvers used by MATLAB^®^ and NEURON).

In the D1 MSNs this biochemical signaling cascade causes synaptic strengthening via several mechanisms, one of which is the phosphorylation of AMPA receptors. As mentioned above, the model instead includes a generic substrate whose level of phosphorylation is the output of the cascade. For the purpose of illustrating the workflow with a proof-of-concept example, we have here linked the fraction of phosphorylated substrate to the AMPA receptor conductance (the conductance is scaled by 1+*f* (where *f*: fraction of the phosphorylated substrate), i.e. when there is very little substrate phosphorylation, very little change in the AMPA conductance is elicited, and vice versa. Modifying the model to include the reactions for AMPA receptor phosphorylation will be made in a future study.

## Discussion

In order to address the growing need for interoperability in biochemical pathway modeling within the neuroscience field, we have developed a workflow that can be used to refine models in all phases of development, keeping in mind the fact that many of the users (including us) are scientists and not professional programmers. For the model and data storage we have chosen the SBtab format which can be easily read and modified by both modelers and experimentalists, and can be converted into other formats, e.g. SBML, MATLAB^®^ SimBiology^®^ or MOD. Our workflow is modularized into different steps allowing the use of each step depending on the need and ensuring interoperability with other tools, such as those described in a similar endeavor named FindSim (Viswan et al. 2018). There are distinct advantages to the workflow, by enforcing a common standard for information exchange, it inherently makes the models more generalizable and reduces the likelihood that simulation results are artifacts of a particular simulator, and nonetheless, it gives users the flexibility to leverage the strengths of each different simulation environment and provides distinct stages of processing that would not be possible in any single simulator. The presented workflow aims to use software components that are free (apart from MATLAB^®^) and solve incremental sub-tasks within the workflow (with open standard intermediate files) to make the workflow easy to branch into scenarios we have not previously considered. It is also possible to circumvent MATLAB^®^ entirely, if desired (e.g. conversion from SBtab to a MOD file that is then used by NEURON).

When deciding which software packages to use we find that an important aspect that must be considered is the cost and licensing. For some researchers, price may be a relevant concern, in other cases a researcher may have to undergo considerable overhead to make their institution/lab purchase a license and possibly operate a license server. Other than MATLAB^®^, we made the choice to disregard commercial products, keeping in line with the field’s trend towards *open source* platforms. We will also expand our tools to support the use of models with more complex geometries, with several compartments (which is also an SBML feature), and tetrahedral meshes that can be used with STEPS in the subcellular simulation setup application. Such advanced geometries can in principle be defined within SBtab tables. When it comes to multiscale simulations, there is the possibility of using the Reaction-Diffusion module (RXD) in NEURON. Currently, however, it does not support the import of SBML as it lacks the concept of spatially extended models, but SBML support might be added to the future versions of RXD (McDougal et al. 2013).

Another interesting consideration is whether the user wants to define any model directly using rules (as in rule-based modeling). This would make the use of Atomizer unnecessary. The transformation from rules to classical reactions seems easier than the reverse, so even if a rule-based simulation is not necessary or too slow, a rule-based description may be shorter and more fundamental in terms of model translation.

When it comes to model analysis, we have here implemented a functionality for global sensitivity analysis. This is a thorough, but computationally demanding approach and the analysis usually needs to be run in parallel on a high performance computing environment. There are also faster but more approximate screening methods that could have been used (Saltelli 2004). In the future we also intend to incorporate uncertainty quantification into the workflow (Eriksson et al. 2019).

While the current workflow is standalone, there are potential alignments to other systems that could assist adoption by the community. For example, other systems such as Galaxy are more general in scope focusing on bioinformatics and are aimed at naive users (Afgan et al. 2016). The current workflow focuses on neuroscience and assumes experienced users. While the current version requires user installation of dependencies, in the future Dockerised versions could potentially help ease installation requirements. Similarly, the Common Workflow Language (Amstutz et al. 2016) is another recent development that could facilitate the exchange of workflow information.

In summary, we have here presented a workflow for biochemical pathway modeling and provided a concrete use case of reward dependent synaptic plasticity, with two additional examples in the supplementary material and the workflow repository. Multiscale models are crucial when trying to understand the brain using modeling and simulations, e.g. how network activity shapes synaptic plasticity or how neuromodulation might affect cellular excitability on sub second timescales. Structured approaches for bridging from detailed cellular level neuron models, to more simplified or abstract cellular-, network-, and even brain region models are developing (Amsalem et al. 2020; Carlu et al. 2020; Schmutz et al. 2020). In the currently illustrated workflow, we add to these efforts by bridging from the subcellular scale to the cellular level scale. We updated parts of the use case model to accomplish a model with only bimolecular reactions that are easier to represent in standards such as SBML. The idea is that users can look at our model as a concrete test case, rerun the workflow (or parts thereof) and then replace the current example model with their own models. In this particular use case, we specifically focused on creating scripts to achieve interoperability between human readable model specification standards and machine-readable standards, and we also wanted to facilitate how a subcellular signaling model could be implemented in different solvers with different strengths, in this case both SimBiology^®^ in MATLAB^®^, as well as STEPS and NEURON. NEURON is currently the most used simulation software for detailed cellular level neuron models, and STEPS can, as said, simulate signaling cascades in arbitrary dendritic morphologies both in a deterministic and stochastic manner. MATLAB^®^ on the other hand has many functions, for example for parameter estimation and we included an implementation for global sensitivity analysis. Several other software is, however, used within the computational neuroscience community for cellular or subcellular model simulations (Ray and Bhalla 2008; Oliveira et al. 2010; Resasco et al. 2012; Akar et al. 2019). To successively make as many of those tools interoperable with standards for both model and data specification, various parameter estimation and model analysis methods, visualization software, etc., will further facilitate the creation of FAIR multiscale modeling pipelines in the future.

## Declarations

### Ethics approval

not applicable

### Consent to participate

not applicable

### Consent for publication

not applicable

### Availability of data and material

see Information Sharing Statement

### Code availability

see Information Sharing Statement

## Acknowledgements

We thank Pavlo Getta for engineering support. We also thank Geir Halnes for sharing scripts for global sensitivity analysis with us. We also thank Sahil Moza for helping with the FindSim use case. The simulations were partly performed on resources provided by the Swedish National Infrastructure for Computing (SNIC) at Lunarc.

## Supplementary Materials

We tested the Subcellular Workflow by running additional models of varying degree of complexity through our tools and various simulation environments discussed in the main manuscript. We imported a pre-existing model of the MAPK cascade (reactions from the green box from [Fig. 7A] from the FindSim workflow in Viswan et al. 2018)18. The model has 102 species, 102 reactions and 160 parameters. We rebuilt the model in SBtab and reproduced simulations of the two test experiments from Viswan et al. 2018 [Fig. 7B - C]. We used 0.1 and 0.001 μmol/l epidermal growth factor (EGF) step pulses as illustrated in Supplementary Fig. 1A and C, respectively, as an input, and phosphorylated MAPK species as output. We performed parameter optimization in MATLAB^®^ as described in the Parameter Estimation section (see main text) on 29 parameters that represent reactions involved in MAPK phosphorylation. Simulated output curves in MATLAB^®^ and COPASI simulations with both original (blue lines) and estimated parameters (red lines) as well as well as the data points used in the model building are plotted in Supplementary Fig. 1B and D. Simulations in BioNetGen needed modifications that included transformation of inositol triphosphate (IP3) producing reactions to mass kinetic form and adding reactions for active protein kinase C (PKC) production instead of the corresponding SBtab expression. The conversion from SBtab to SBML and from SBML to BNGL models was done by our conversion tools (see paragraph titled Simulations in STEPS). The web-based subcellular simulation setup application was used for importing of the BNGL model to STEPS and running simulations. Note that both stochastic solvers we used produce qualitatively similar solutions which does not converge to deterministic solutions for the same model. This result is expected for a general type nonlinear and highly stochastic model. In simulations in NEURON output of the cascade has been coupled to affect the conductance of Kv4.2 channels in the dendrites, as if a global EGF signal had arrived to the whole neuron. MAPK can phosphorylate Kv4.2 channels which decreases their conductance, and this effect is implemented by scaling the maximal channel conductance. Phosphorylation of the Kv4.2 channels in the dendrites makes the neuron more excitable, which starts firing due to random synaptic input distributed across the dendrites (Supplementary Fig. 5C). Good agreement was found between simulations performed in MATLAB^®^ (Supplementary Fig. 2), COPASI (Supplementary Fig. 3), STEPS, BioNetGen (Supplementary Fig. 4) and NEURON (Supplementary Fig. 5). We also performed a global sensitivity analysis (see Global Sensitivity Analysis in the main text) on the estimated parameters to see what effect different parameters have on the output, either by themselves or in interaction with other parameters (Supplementary Fig. 3). Our SBtab files and corresponding conversions are linked in the main repository for this manuscript, alongside the example used in the main text.

Finally, we rebuilt a model of the EGF-dependent protein kinase B (Akt) pathway in SBtab (Fujita et al. 2010 provided by Hass et al. 2019 as one of the benchmark problems for modeling intracellular processes), with 11 species, and performed a global sensitivity analysis on the parameters and the figures are available in the Subcellular Workflow GitHub repository19.

**Supplementary Fig. 1:**
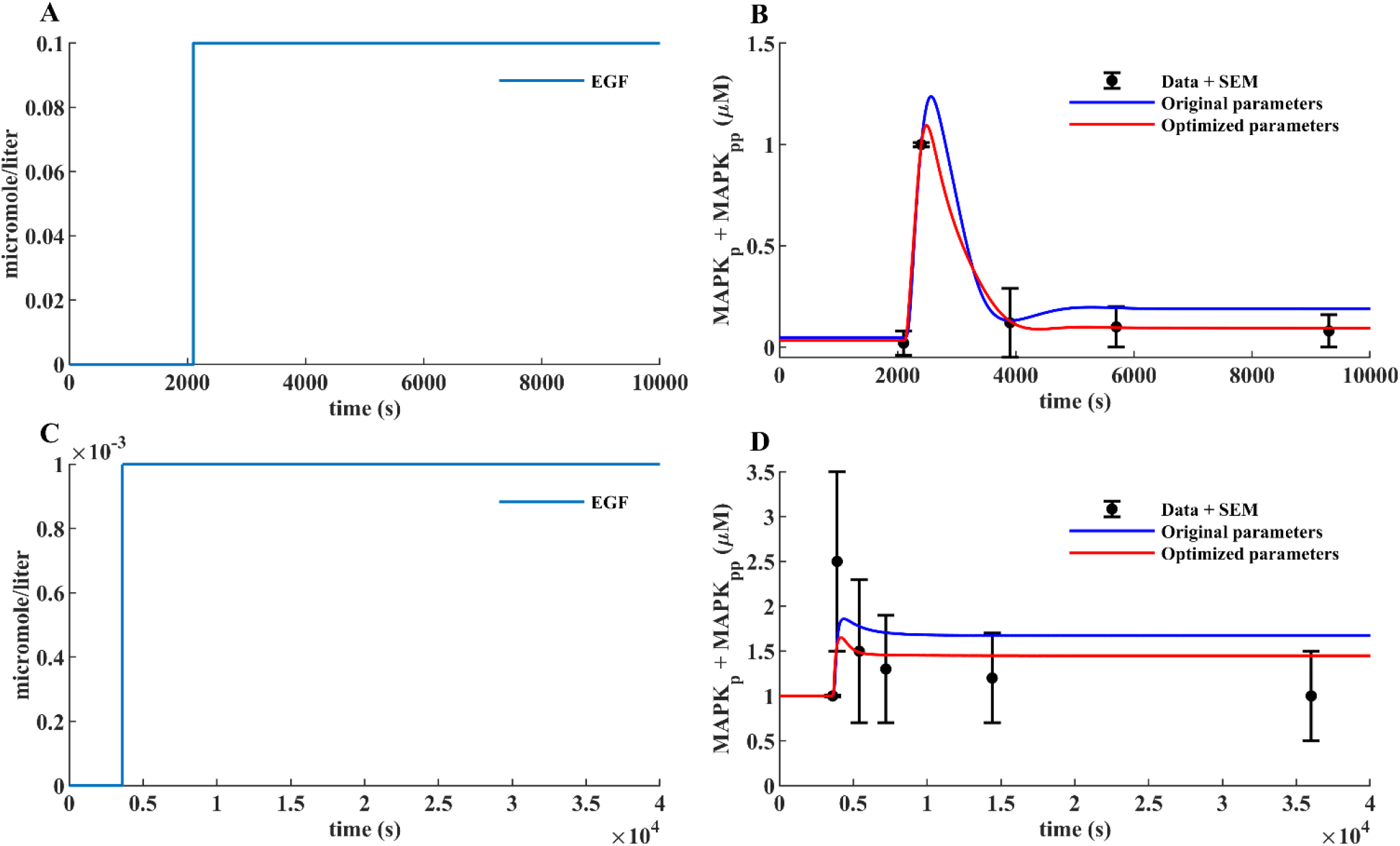
MATLAB^®^ simulations of the MAPK cascade from the example model provided by FindSim (Viswan et al. 2018). (A, B) and (C, D) correspond to figures 7b and 7c in their manuscript, and to experiments E0 and E1 in our files. (A) and (C) show inputs for the two experiments. Step input of EGF at 0.1 and 0.001 μmol/l is used. (B) and (D) show measured output consisting of the sum of phosphorylated MAPK species. The black dots and error bars refer to data points and standard error obtained from the publication; the blue lines represent model behavior when simulating with the original parameters; the red lines represent model behavior when simulating with parameters obtained by optimization using our MATLAB^®^ parameter estimation tools. The plots are analogous to simulations in Fig. 3.

**Supplementary Fig. 2:**
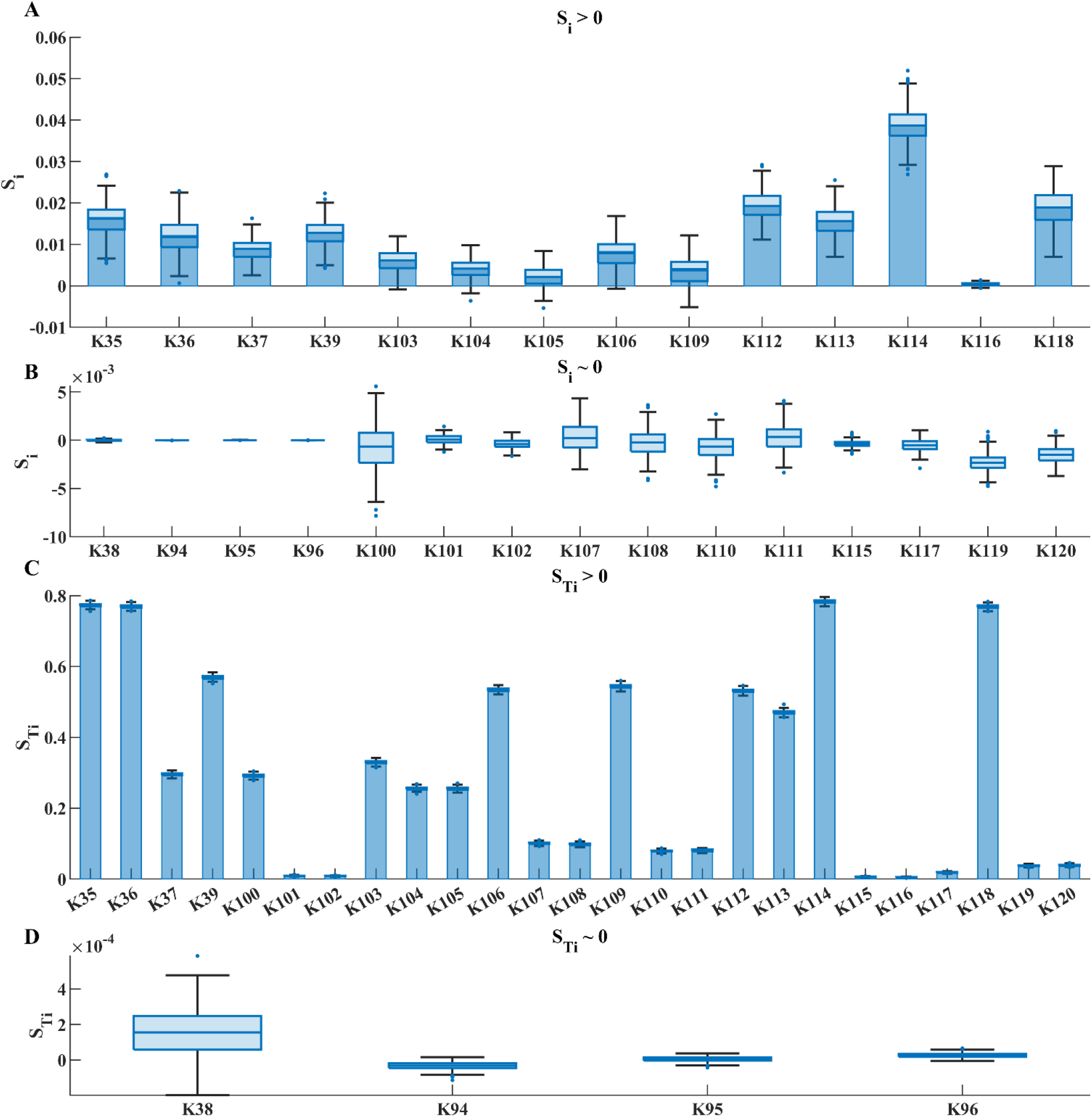
Bar graphs of the first order, Si, and total order, STi, sensitivities indices of the 29 parameters that represent reactions involved in MAPK phosphorylation for one of the experiments (E0) are shown (panel A-B and C-D, respectively). The sensitivities indices are defined in the main text and were calculated based on the scores used in the optimization for the experiment. The parameters, Θ, were sampled independently from a multivariate lognormal distribution with log10(Θ)~ N(μ, σ), using μ =log10(Θ*) and σ=0.1, where Θ* correspond to the optimal values received from the optimization. A sample size of N=100000 was used (corresponding to 3100000 reshuffled samples used in the calculations (Saltelli 2004)). The analysis took approximately 8 hours on 18 compute cores (Intel 10980K). The uncertainty in the sensitivity indices due to sampling was estimated through bootstrapping and indicated by boxplots, showing the median, 25- and 75-percentile, the distance between the lower and upper quartile is the interquartile range (IQR), values that are more than 1.5 times the IQR distant from the top or bottom of the box are considered outliers and all other values are included in the whiskers of the boxplot.

**Supplementary Fig. 3:**
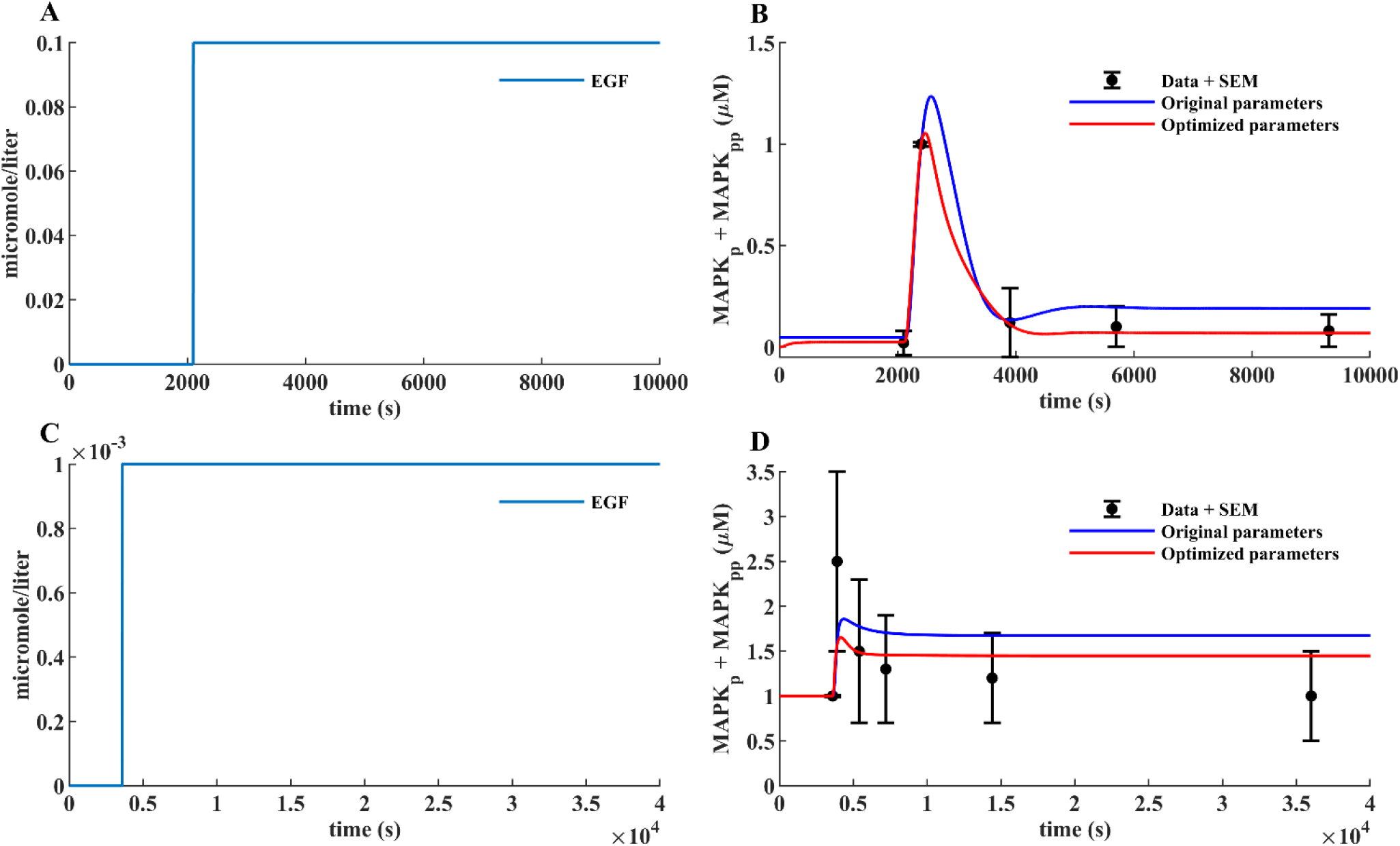
COPASI simulations of the MAPK cascade from the example model provided by FindSim (Viswan et al. 2018). (A, B) and (C, D) correspond to figures 7b and 7c in their manuscript, and to experiments E0 and E1 in our files. (A) and (C) show inputs for the two experiments. Step input of EGF at 0.1 and 0.001 μmol/l is used. (B) and (D) show measured output consisting of the sum of phosphorylated MAPK species. The black dots and error bars refer to data points and standard error obtained from the publication; the blue lines represent model behavior when simulating with the original parameters; the red lines represent model behavior when simulating with parameters obtained by optimization using our MATLAB^®^ parameter estimation tools. The plots are analogous to simulations in Fig. 5

**Supplementary Fig. 4:**
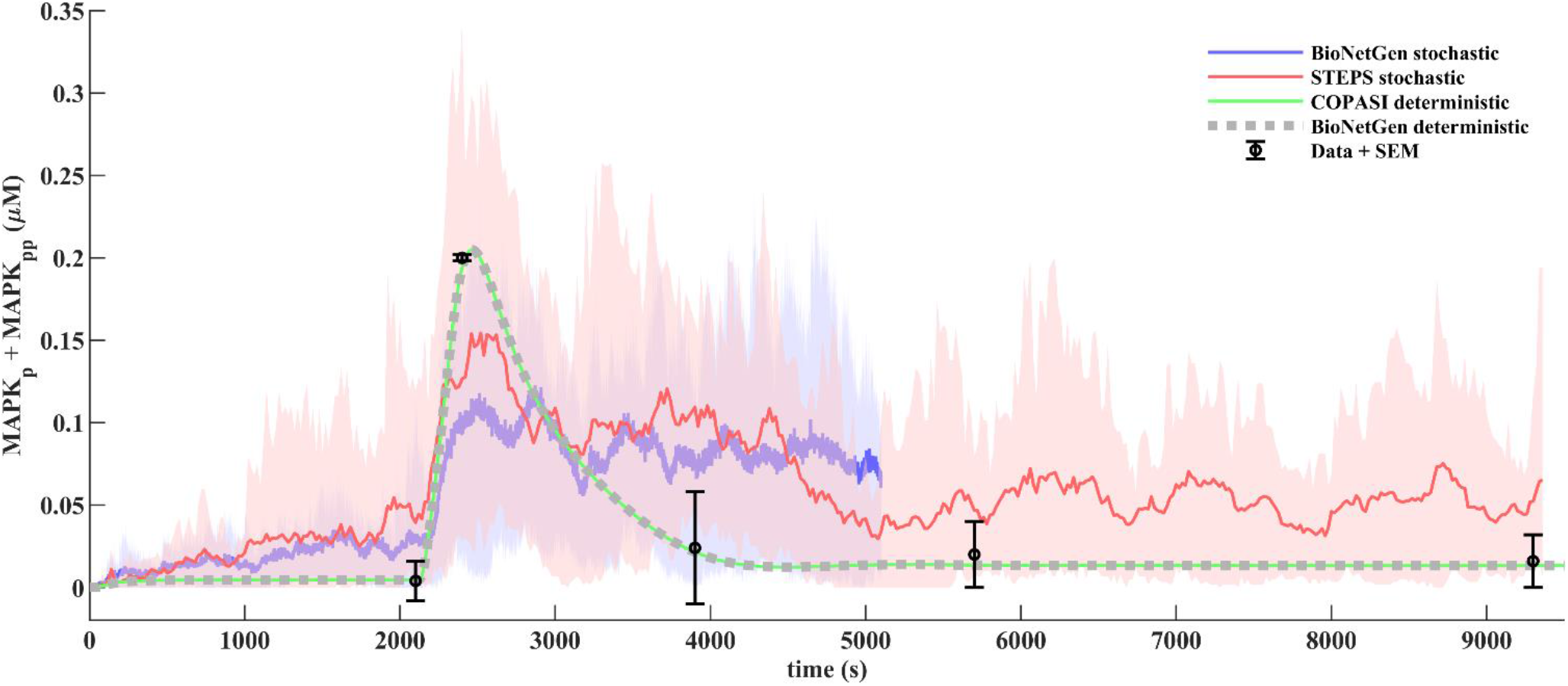
STEPS and BioNetGen simulations of the MAPK cascade from the example model provided by FindSim (Viswan et al. 2018). The optimized parameter set obtained using our MATLAB^®^ parameter estimation tools was used for the simulations. Data and simulations correspond to figure 7b in Viswan et al. 2018, to experiment E0 in our files and to red lines on Supplementary Fig 1 and 3. Step input of EGF at 0.1 μmol/l was applied starting from 2100 second of simulation. Simulated traces show measured output consisting of the sum of phosphorylated MAPK species. The black dots and error bars refer to data points and standard error obtained from the publication; the dotted green line represent model behavior when simulating with COPASI deterministic LSODA solver; the magenta line was obtained by deterministic simulation of BioNetGen solver obtained by the automatic conversion of the example SBtab model modified to make it compatible with BioNetGen language. Note the close correspondence of optimized example model and modified BNGL model solutions; the blue curve corresponds to BioNetGen stochastic ssa solver simulation of the modified BNGL model averaged by 15 simulated traces (5000 seconds of simulations are shown). The blue colored area represents 10%-90% confidence interval of the solution; the red line represents STEPS stochastic solver simulation of the modified BNGL model averaged by 35 simulated traces. The red colored area represents 10%-90% confidence interval of the STEPS solution.

**Supplementary Fig. 5:**
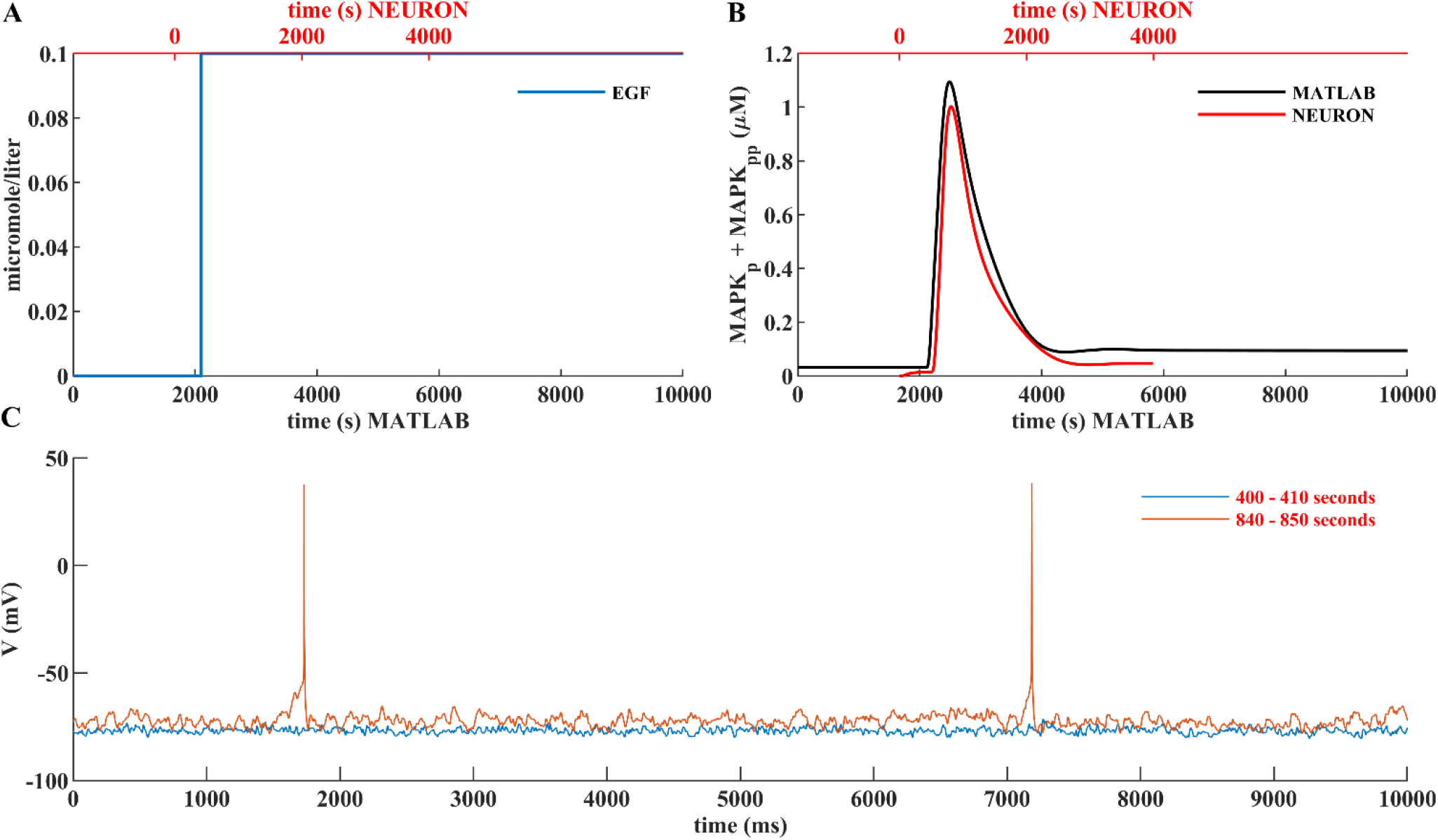
NEURON simulation of the MAPK cascade. The output of the cascade has been coupled to affect the conductance of Kv4.2 channels in the dendrites, as if a global EGF signal had arrived to the whole neuron. MAPK can phosphorylate Kv4.2 channels which decreases their conductance, and this effect is implemented in by scaling the maximal channel conductance according to g = (1 − ([MAPKP] + [MAPKPP])/[MAPK]total) gmax. (A) The simulation lasts 4000 s, and the EGF input arrives at 500 s. (B) Comparison of the normalized output of the cascade in NEURON and MATLAB^®^. (C) 10s-long somatic voltage traces before (at 400 s) and after (at 840 s) the EGF stimulus show the effect of the cascade on somatic voltage.

## Footnotes

1 for example: at the *time of writing* the SimBiology^®^ toolbox (in MATLAB^®^) exports the *time* variable as a normal variable, without the required definition URL, it must be added manually.

2 Kernel density estimation does not work very well in high dimensional problems.

3 A library providing an application programming interface for SBML.

4 https://sbtab.net/

5 https://www.sbtab.net/sbtab,/default/converter.html

6 https://subcellular-workflow.readthedocs.io/

7 An example of the settings file can be found in the GitHub repository https://github.com/jpgsantos/Subcellular_workflow/

8 https://subcellular-workflow.readthedocs.io/

9 Where *V* is the variance operator and *E* (conditional) expected value.

10 Other distributions can be used as well.

11 Pseudo random number generators have a *seed* parameter for initialization.

12 Conversion function available in https://github.com/jpgsantos/Subcellular_workflow/ and instructions in https://subcellular-workflow.readthedocs.io/

13 Including, but not limited to: Python (NumPy), C (GNU Scientific Library (GSL), CVODE), GNU Octave (LSODE), R (deSolve), and core MATLAB^®^ (e.g. for ode15s).

14 It should be noted that the units may not show up correctly in COPASI (depending on the version) even if they are correct in the SBML file itself.

15 The online application for subcellular simulations can be found in https://subcellular.humanbrainproject.eu/model/simulations with the documentation in https://humanbrainproject.github.io/hbp-sp6-guidebook/online_usecases/subcellular_level/subcellular_app/subcellular_app.html

16 A file given either as a series of tab separated text files or one open document spreadsheet file. Some content of the SBtab model is mandatory, some optional.

17 libSBML must installed with R bindings: i.e. SBML output is optional.

18 https://github.com/jpgsantos/Model_Viswan_2018

19 The model GitHub can be found in https://github.com/jpgsantos/Model_Fujita_2010 and example plots of the GSA are in https://github.com/jpgsantos/Model_Fujita_2010/tree/master/Results/Examples

